# High Resolution Multi-Pass Astral Analyzer Quantification Enables Highly Multiplexed 35-Plex Tandem Mass Tag Proteomics

**DOI:** 10.64898/2026.02.24.707764

**Authors:** Hamish Stewart, Steven R. Shuken, Christopher Rathje, Julia Krägenbring, Martin Zeller, Tabiwang N. Arrey, Bernd Hagedorn, Eduard Denisov, Robert Ostermann, Dmitry Grinfeld, Johannes Petzoldt, Daniel Mourad, Philipp Cochems, Florian Bonn, Bernard Delanghe, Michael Wiedemeyer, Alexander Wagner, Ryan Bomgarden, Dustin C. Frost, Nathan R. Zuniga, Ramin Rad, João A. Paulo, Eugen Damoc, Alexander Makarov, Vlad Zabrouskov, Christian Hock, Steven P. Gygi

## Abstract

Tandem mass tags (TMT) allow highly multiplexed and thus high-throughput, precisely quantitative proteomic analysis. Incorporation of additional deuterated reporter channels has near-doubled the multiplexation achieved with Thermo Scientific™ TMTpro reagents from 18 to 35-plex but requires extremely high ∼100k analyzer resolving power at m/z 128 to differentiate and quantify reporter ion channels, far beyond any single reflection time-of-flight analyzer, and exceeding the multi-reflection Thermo Scientific™ Astral™ analyzer in its standard operation. A multi-pass mode of Astral operation has been developed for the Thermo Scientific™ Orbitrap™ Astral™ Zoom mass spectrometer that triples the ion path to 90 m, more than doubling resolving power for a narrow m/z range. This “TMT HR mode” has been integrated into a new method of TMT proteomic analysis that splits regular MS2 analysis of labeled peptides into paired measurements comprising wide mass range scans for peptide identification, and TMT HR mode scans for reporter ion quantification. The method has been shown to accurately quantify 32-plex labeled HeLa protein lysate and provide far greater depth of analysis as state-of-the-art Orbitrap-only methods, while analysis of 11-plex labeled yeast showed no analytical depth sacrificed vs regular Orbitrap Astral TMT analysis. Further comparative measurements of a 2-cell line 35-plex sample demonstrated greater analytical depth, and similar quantitative precision, to “gold standard” Orbitrap MS3 methods.

## INTRODUCTION

Isobaric chemical tags, such as tandem mass tags (TMT), remain a valuable tool in mass spectrometry-based peptide quantitation^1^. Samples tagged with different isobaric labels generate isotopically differentiated, and thus m/z differentiated, reporter ions upon collisional fragmentation, allowing individual quantitation of multiplexed samples without increasing the complexity of the unfragmented mass spectrum. The early 6-plex tags expanded to 18-plex, many reporter ions separated only by the 6 mDa mass difference of a ^13^C to ^15^N substitution, requiring relatively high >50k mass resolution at m/z 128 to isolate^2,3^. This demand is incompatible with contemporary single-reflection time-of-flight mass analyzers, as even sub-nanosecond detector response times cause excess broadening of measured 10s of µs-level ion flight times.

Multi-reflection analyzers, such as the Astral (ASymmetric TRAck Lossless) analyzer, benefit from an order-of-magnitude longer ion path, typically managing 70k, and at least 55k, resolution at m/z 128, but falling to below 50k under the space charge influence of 1000 ions in a single m/z peak^4^. The Thermo Scientific™ Orbitrap™ Astral™ mass spectrometer^5,6^, which incorporates both Astral and Orbitrap analyzers, has previously been shown to be highly advantageous in the analysis of such highly multiplexed TMT-labeled samples^7-9^.

The introduction of 35-plex TMT^10^, which incorporates deuterated tags, manifest reporter ions in quadruplets with 3 mDa spacing, severely challenging mass analyzers with a requirement for approximately 100k resolving power at low m/z. For the Orbitrap Astral mass spectrometer, this is simply incompatible with the performance-driving fast and sensitive Astral analyzer, at least in this incarnation and in normal operation.

For multi-reflection analyzers including the Astral analyzer, it has long been shown that a multi-pass mode, trapping ions within to make additional passes of the multi-reflection flight path, may greatly enhance achievable resolving power^11-13^. Figure 1 illustrates the operation within the Astral analyzer, whereby ions are injected from an ion trap source, and deflected into the analyzer by a first and then a second prism deflector, the Relay Prism, which sets the 2.2-degree injection angle into the multi-reflection path. While passing through the multi-reflection path, this prism deflector is set to a trapping mode, and ions are reflected back to make 3 full passes before it is set back to a transmitting mode and extracts ions to the detector. As resolving power is dependent on the length of the ion flight path, the 3x longer path achieved by the multi-pass method enhances peak resolution substantially^13^.

**Figure 1.**
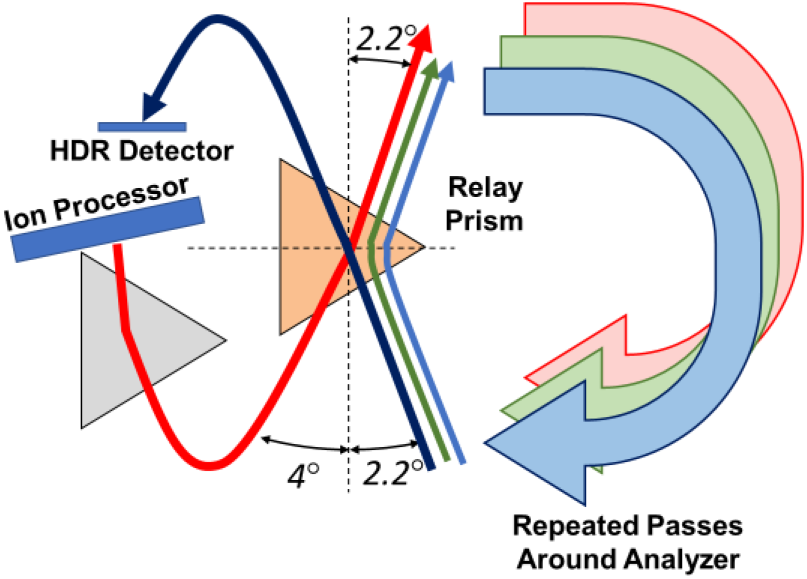
Illustration of ion path through the prism deflector during multi-pass operation, from injection (red), internal reflections (green then blue) to extraction of ions to the detector (purple).

This method is not directly compatible with TMT analysis, however, as it induces a reduction in mass range. Light ions lap heavy ions and the number of passes an ion makes before reaching the detector becomes ambiguous, unless the ion m/z is known. While TMT reporter ion m/z *is* known, peptide fragment ions required for peptide identification are not, thus terminating the ability to simultaneously monitor peptide fragments and TMT reporter ions. Concepts exist to deconvolute such spectra over multiple injections^13^ but have not been properly realized.

A new data-dependent method of TMT-labeled peptide analysis has been developed for an improved Orbitrap Astral instrument, the Orbitrap Astral Zoom mass spectrometer, whereby the Orbitrap analyzer acquires full-MS survey scans as normal, while the Astral analyzer’s data dependent MS2 acquisition is split into two stages. The first is a regular MS2 analysis to monitor peptide fragments, followed by a second acquisition in high-resolution multi-pass “TMT HR” mode to quantify TMT reporter ions only^14^. Suitably tuned, this method provides adequate resolving power to separate 35-plex TMT reporter ions at up to 500-1000 ions in a multiplet.

## EXPERIMENTAL METHODS

### Instrumentation

The layout of the Orbitrap Astral Zoom mass spectrometer is shown in Figure 2. It incorporates an electrospray ion source and quadrupole mass filter, along with Orbitrap and multi-reflection Astral mass analyzers. The instrument’s structure and operation have been broadly described^5,6^, whereby electrosprayed ions are accumulated in the IRM (Ion Routing Multipole) and then returned to the C-Trap and injected into the Orbitrap analyzer for full-MS survey measurement. For MS2 analysis, isolated analyte ions are passed into the IRM, and then into the Ion Processor for collisional dissociation and cooling, followed by pulsed extraction into the Astral analyzer.

**Figure 2.**
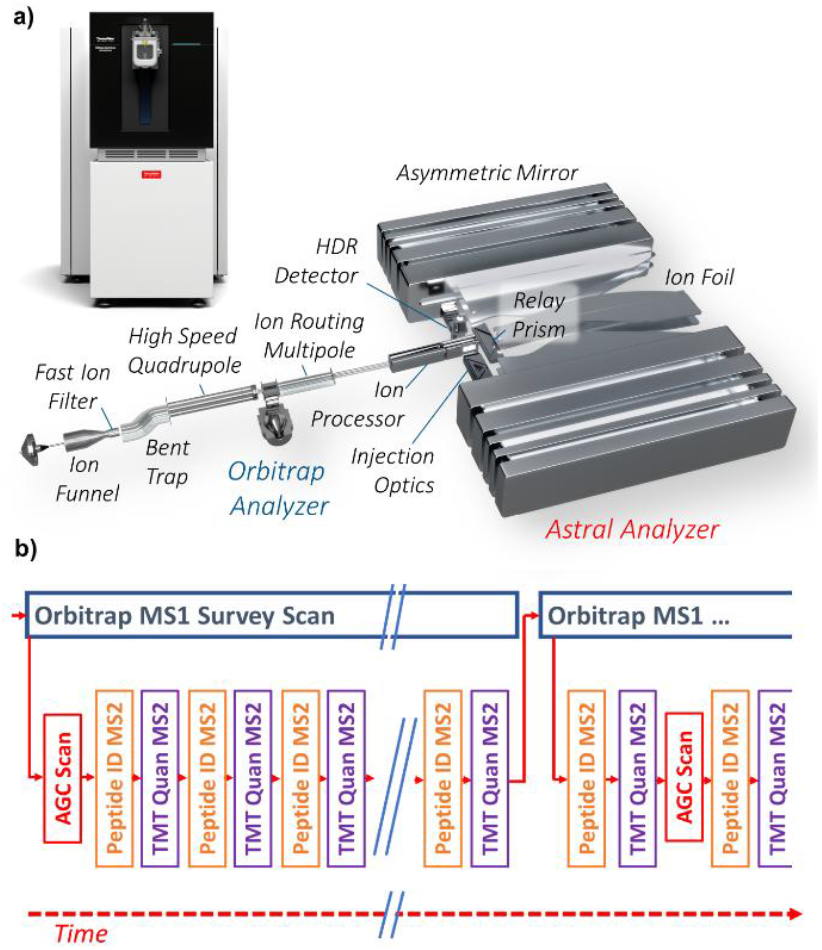
a) External appearance and ion optical layout of the Orbitrap Astral Zoom mass spectrometer. b) Order of scan events in data dependent TMT 32-plex application.

In the new method, labeled peptide precursors are detected by MS1 survey scans and identified in data-dependent MS2 analyses as typical to TMT analysis. However, each of these measurements is supplemented by a second MS2 scan dedicated to quantifying the TMT reporter ions. Isolated tagged-peptide ions are fragmented, albeit with optionally higher normalized collision energy (NCE) to maximize reporter ion yield. However, analysis occurs via multipass mode, in which the Relay Prism (a trans-axial deflector with a wide spatial acceptance^6^) is switched from a transmitting mode (-160 V) to a trapping mode (+250 V), that causes returning ions to be completely reflected and to make a second and a third pass through the multi-reflection analyzer. The Relay Prism is then set back to a transmitting mode before the ions complete the third pass, allowing them to be extracted to the detector. The switching time is as fast as 20 µs.

A significant hurdle of switching between multi-pass and single-pass modes of operation is that the carefully calibrated focal plane position of the ion packet becomes misaligned from the detector surface, destroying time-resolution. Normally the alignment is performed by tuning voltages of the mirror electrodes prior to operation. The tuning process is, however, slow because the power supplies incorporate large capacitors to filter electronic noise for stability, and the high voltages require almost 2 seconds to settle after any modification. Such a long dead time is incompatible with a scan-to-scan cadence of 100 Hz or higher.

An alternative method for rapidly making small shifts to the focal plane position is to adjust the average velocity of ions in flight by means of a low-voltage compensation electrode similar to the existing Ion Foil electrodes mounted between the mirrors, but lacking any specific shape function^15^. This method performs a first-order adjustment of the time focus without any need to adjust high voltages.

Lacking space for a new electrode, it was at first thought that the entire assembly of inter-mirror electrodes and their normally grounded mountings could be floated to this correction voltage, however the details of the analyzer’s design made that rather challenging. A simpler alternative was realized, in that the first electrode of each ion mirror, being a long straight electrode and normally grounded, could serve instead! This was thus floated and connected to a low-voltage supply with a relatively fast sub-ms switching regulator. Switching this parameter to around -30 V is sufficient to correct the focal plane position (Supplemental Figure 1), though variation is substantial between different instruments.

This voltage supply can also improve robustness to space charge, as a major effect of resonant space charge on a peak is to shift the focal plane position proportionally to the number of charges in this peak^16^. A small compensation voltage may reverse this effect. To enhance the dynamic range, the focal plane position is kept slightly overtuned which compensates for space-charge-induced shifts for peaks of greater intensities^4^. For TMT reporter ions, the impact of space charge is the limiting factor of dynamic range; resolution falls under increasing ion number until the ability of the peak deconvolution algorithm^17^ to separate overlapping peaks is defeated. TMTpro 32-plex reporter ions appear as quadruplets with 3 mDa mass separation, and as space charge effects carry over between similar m/z species, the effective ion capacity per multiplet is split between four species, far more challenging compared to acquisition of TMTpro 18-plex or TMT 11-plex samples, which produce only reporter ion doublets.

Worse, there is a ∼40-65% allowed sacrifice of ion transmission associated with the multi-pass mode, thus again doubling the number of ions injected into the analyzer for a given number of ions detected, effectively again doubling the required space charge tolerance.

These effects challenge the viability of the method. Fortunately, the TMT HR mode of operation is dedicated solely to measurement of known m/z ions, so the highest achievable resolving power may be sacrificed for space charge tolerance. Figure 3 shows resolving power of m/z 138 ions generated from Pierce™ FlexMix™ calibration solution, at different numbers of ions in a peak controlled by varying the Ion Processor’s ion accumulation time. It may be seen that resolving power greatly exceeds the 90k level used in Orbitrap TMT32-plex analysis at 500 ions detected (while the number of injected ions was likely 1000 under the 40-65% transmittance). This is a stark contrast to the standard mode’s calibration optimum, that returns higher resolving power at low ion number, where it is normally desirable for good statistical mass accuracy^4^.

**Figure 3.**
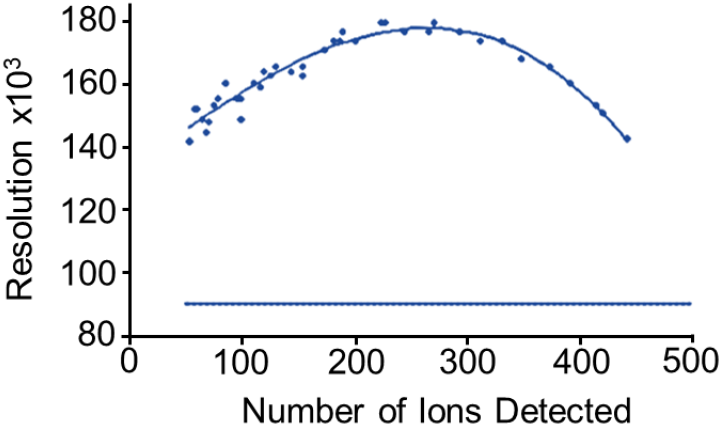
Example of high space-charge tolerance tuned system, showing FlexMix m/z 138 resolving power vs number of ions detected (∼50% transmission with respect to the standard multi turn mode).

### Evaluation Experiments

Proof of concept measurement of TMT HR mode’s ability to separate closely spaced reporter ions was made via measurement of an infused mixture of TMTpro and deuterated TMTproD reagents, to reproduce a single reporter ion quadruplet. A practical evaluation of the method, however, required complex TMT labeled samples.

Pierce TMT11plex yeast digest standard (TKO) was used for method optimization and as an evaluation standard. There is currently no such manufactured standard for TMTpro 32- or 35-plex experiments, so instead a TMTpro 32-plex HeLa lysate digest test sample was made with defined 1:4 concentration ratios between different channels, shown in Figure 4a. Here, most of the deuterated tags are present at fourfold higher concentration than their non-deuterated neighbors, creating more challenging reporter ion multiplets in which high- and low-intensity reporter ions are adjacent. This arrangement maximizes susceptibility to interference from crosstalk when closely spaced peaks are not fully resolved.

**Figure 4.**
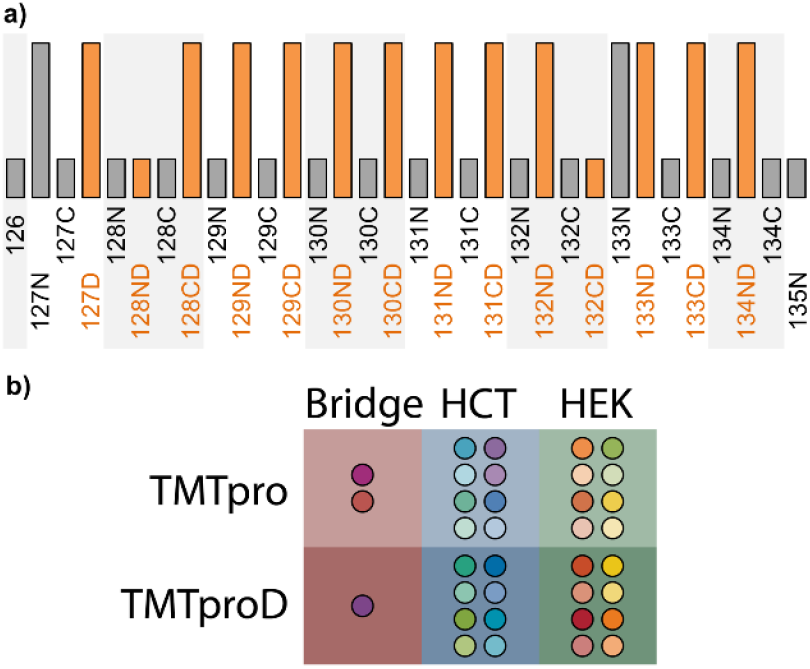
a) Relative ratios of multiplexed TMTpro labeled HeLa to produce known 1:4 quantitative ratios within reporter ion multiplets. b) Labelling of HCT, HEK, and mixed bridging samples to create a 2-cell line 35-plex test sample.

An additional test of quantitative performance used multiplexed lysate digests from two human cell lines (HCT116 and HEK293T) with eight replicates of each labeled with TMTpro and TMTproD reagents. The remaining three channels to reach the 35-plex capacity were allocated to bridging samples: mixtures of equal quantities of the 16 primary samples. These bridges were labeled and used in data processing to correct for the retention time shift between deuterated and non-deuterated labeled samples. The occupation of samples to the different TMTpro channels is summarized in Figure 4b, similar to prior work^10^ albeit without fractionation.

### LCMS Methods

Differing amounts of sample were injected by a Thermo Scientific™ Vanquish™ Neo LC system configured for direct injection and separated on an IonOpticks™ Aurora Frontier™ XT (75 µm x 60 cm) column with either 78 or 88-minute gradients (80 and 90-minute injection cadence). A field asymmetric ion mobility (FAIMS) device, the Thermo Scientific™ FAIMS Pro Duo interface, was used in some methods to filter electrosprayed ions prior to admission into the mass spectrometer, reducing chemical interferences and typically improving quantitative performance in TMT analysis^18^.

The instrument was operated in a Top20 data dependent analysis (DDA) mode, with a 120k Orbitrap MS1 survey scan of m/z 400-1500 and a 300% AGC target (∼300,000 ions). Peptide identification Astral MS2 scans were made with a 1.2 m/z isolation window and 5 ms max inject time, Normalized collision energy (NCE) 32%, 150-2000 m/z range and 100% AGC target (10,000 ions). By contrast, the TMT HR mode quantification scans were set with a more finely isolating 0.5 m/z window, to minimize interferences, a longer 50 ms max ion accumulation time to cover for ion losses associated with fine isolation, a 300% AGC target and a higher 55% NCE, which favors reporter ion generation. The ability to separately optimize experimental settings for both peptide identification and reporter ion quantification is a compensatory factor to the cost of needing two scans for every precursor.

Additionally, the instrument software allows a high sensitivity Low Input mode, in which the gain of the detector is raised to increase signal/noise and probability of single ion detection. Normally, this is intended for very low load or single cell applications^19^ as higher signal area presents a higher risk of detector saturation and poor quantitation. In this case, however, the long ion travel times, relatively broad peaks, and low intensity of individual reporter ions measured via TMT HR mode were considered to reduce the risk. Low input mode was activated for a comparative test with the 1:4 TMT 32-plex Hela sample.

For comparison to state-of-the-art TMT analysis, duplicate measurements of 250 ng of the 2-cell line TMTpro 35-plex sample were carried out on the Orbitrap Astral Zoom mass spectrometer operating in a standard DDA mode, without TMT HR quantification scans, and a Thermo Scientific™ Orbitrap™ Eclipse operating with real time search, synchronous precursor selection MS3 TMT quantification^20,21^.

Data Processing: Data processing was performed with a beta version of Thermo Scientific™ Proteome Discoverer™ 3.3 software with options designed for TMT HR mode analysis, as well as capability to filter TMT reporter ion peaks by signal to noise ratio and resolution, particularly useful for removing space charge saturated, unresolved quadruplets. Results were searched using the SEQUEST HT algorithm and precursor detector node, as well as the INFERYS® rescoring node. All results were filtered for protein groups and peptides being quantified in all multiplexed channels.

## RESULTS AND DISCUSSION

Tuning of the TMT HR mode proved surprisingly difficult across different analyzers compared to the original hand-tuned investigation^13^, possibly due to the longer flight paths used in more recent iterations of the Astral analyzer, which made control of ion dispersion more challenging. Nevertheless, it was possible to achieve a viable machine calibration, though transmission suffered significant 40-65% losses. Figure 5 shows a test example of a TMTpro reporter ion quadruplet generated from MS2 acquisition of a TMTpro reagent sample mixed with a 1:10:1:10 ratio. The resolution of the peaks is sufficient for them to be consistently well deconvoluted and identified, and the ratios are consistent, if not quite the intended ratios which most likely comes from the sample itself. Cross checking Orbitrap and Astral reporter ion spectra showed matching ratios in both analyzers. The peak shapes deviate somewhat from the gaussian ideal, with some peak fronting in addition to the usual tailing, that likely would somewhat hinder large deviations between multiplexed sample concentrations.

**Figure 5.**
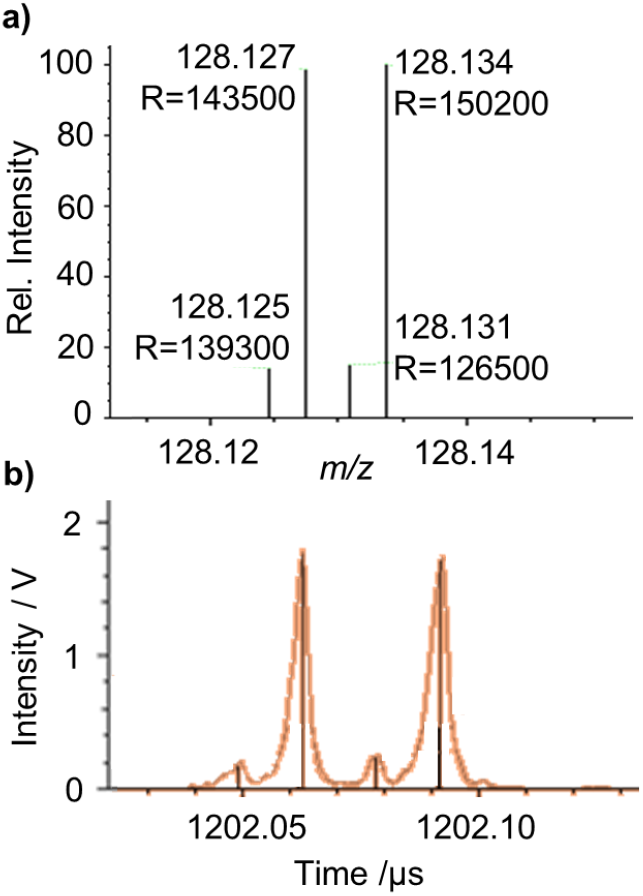
a) mass and b) time-profile spectra of TMTpro reporter quadruplets.

### TKO Yeast and HeLa Benchmarking

The first application test was to how TMT HR mode compared to regular single-pass operation with a common TMT standard, 500 ng Pierce TMT11plex yeast digest, separated over a 78-minute gradient. It was intuited that there might be a major sacrifice in performance with TMT HR mode due to the doubled number of scans per precursor, but in practice a similar number of quantified peptides and protein groups were returned, shown in Figure 6a. Given how short the peptide identification scans are compared to TMT quantitation scans (5 vs 50 ms max inject time), the impact of the two-stage measurement strategy may be minimized, while the higher resolution of the TMT HR scan increases the tolerance of the reporter ion measurement to space charge and saturation effects. The number of quantified protein groups was lower than previous published result,^5^, though FAIMS was not used in these experiments.

**Figure 6.**
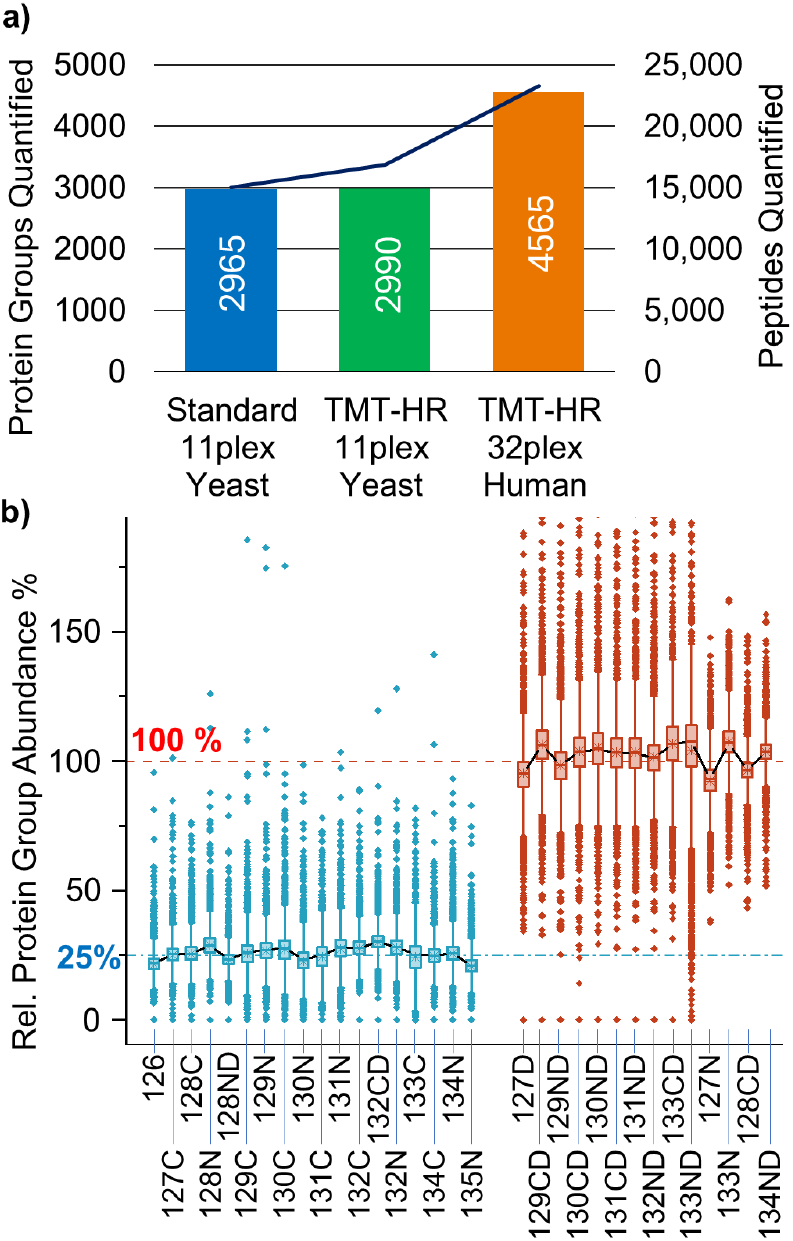
a) Protein groups and peptides identified over 78-minute gradients in standard and TMT HR methods, with 500 ng TMT11plex yeast standard or 1:4 ratio TMTpro 32-plex HeLa sample. b) Relative quantitation of reporter channels in TMT HR measurement of 1:4 ratio TMTpro 32-plex HeLa sample. Boxes show 25-75% spread, and lines double the interquartile range, with outlier points surrounding.

Of the three knockout genes in this standard sample, an average of ∼15% TMT reporter ion signal was measured in the empty channels. This residual signal arises from co-isolation interference by other labeled precursors and is an inherent limitation of MS2-based quantification for quadrupole-isolated precursors, independent of analyzer resolving power.

The same experiment was performed with 500 ng of the 1:4 ratio TMTpro 32-plex labeled HeLa sample, at which point the number of proteins quantified in single pass operation collapsed to 10 (Supplemental Figure 2), as expected because unresolved peaks were excluded during processing. By comparison, >4500 protein groups were quantified with TMT HR mode. The relative abundances of the 32 channels are shown in Figure 6b, and the 1:4 ratios of the two channel groups can be seen to be tightly reproduced, with narrow variation for the majority of points. There was little evidence of severe cross-talk between adjacent peaks that would amplify the apparent signal of low-intensity peaks. Longer gradients and higher loads moderately improved the number of quantified proteins to >5000 (Supplemental Figure 3).

One notable observation is that it was presumed that the ability to individually set isolation range and collision energy for the peptide identification measurements and the TMT HR quantitation scans would prove extremely important. While the two certainly had differing optimal values (NCE 35% for ID vs 55% for TMT, narrow isolation being considered a prerequisite for reducing channel interferences and subsequent ratio compression), the impact during method development was more modest than hoped for, without great effect on the number of reported quantified peptides. This experiment provides good evidence that the TMT HR mode is suitable for TMTpro 32-plex labeled peptide analysis, though it is likely that there is still room to further optimize the methodology, particularly when FAIMS separation is employed.

### Low Input Mode

The impact of low input mode, which raises the detector gain, was also evaluated. 250 ng of a similar 32-plex HeLa sample was acquired over a 90-minute cycle (equivalent to 560 samples per day) with FAIMS separation at 40, 60 and 80 V compensation voltages. Practically, low input mode increases the voltage applied across the detector’s photomultiplier tube by approximately 50 V, and the impact of the resultant higher gain is shown in Figure 7. The median S/N of reporter channels increased from less than 50 to more than 130, shifting the distribution of intensities upwards, and without obvious truncation. With low input mode activated, the number of quantified protein groups increased from 1750 to 2750; a huge improvement, albeit one from a relatively sensitivity-restricted method. In regular mode, a single ion peak yields a signal-to-noise ratio of ∼2, implying that most TMT channels contained only a few tens of ions, with the space-charge restricted upper limit being somewhat more than 100 ions unless spectral averaging is applied, which is rarely used in Orbitrap Astral methods.

**Figure 7.**
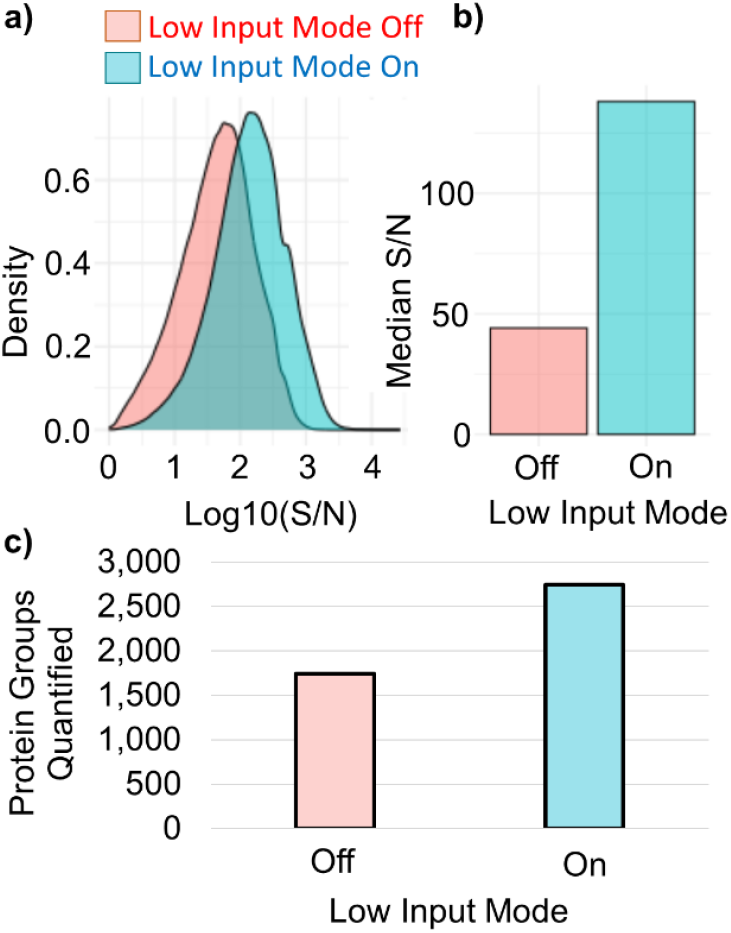
Impact of low input mode on a) signal to noise ratio distribution, b) median S/N and c) quantified protein groups from 78-minute gradient analysis of 250 ng TMTpro 3-plex labeled sample.

This experiment was carried out separately to that reported in Figure 6, with a slightly different method involving use of FAIMS separation, and on a degraded sample which returned substantially lower quantified protein numbers (Figure 7). Nevertheless, low input mode more than doubled the typical signal-to-noise ratio, resulting in a >50% boost in quantified protein groups.

### 2-Cell Line 35-plex

250 ng of the 2-cell line sample was analyzed in a 90-minute full-cycle experiment via both regular Orbitrap Astral Zoom MS2 quantification, TMT HR mode, and a comparative RTS-SPS-MS3 experiment by an Orbitrap Eclipse mass spectrometer. The latter method represents a gold standard for TMT analysis. This time, the regular Astral MS2 quantification returned a number of quantified proteins, perhaps owing to the heterogeneity of the multiplexed samples. The TMT HR mode granted the greatest depth of analysis, at 2320 quantified proteins vs the Orbitrap Eclipse’s 1330, shown in Figure 8. Greater depth is consistent with prior comparisons of Orbitrap vs Orbitrap Astral analysis^5,7^; the Astral analyzer, even with a very long max 50 ms inject time, is still much faster than an Orbitrap analyzer set to 90k resolving power (180 ms, or 90 ms in TurboTMT mode), and several times more sensitive.

**Figure 8.**
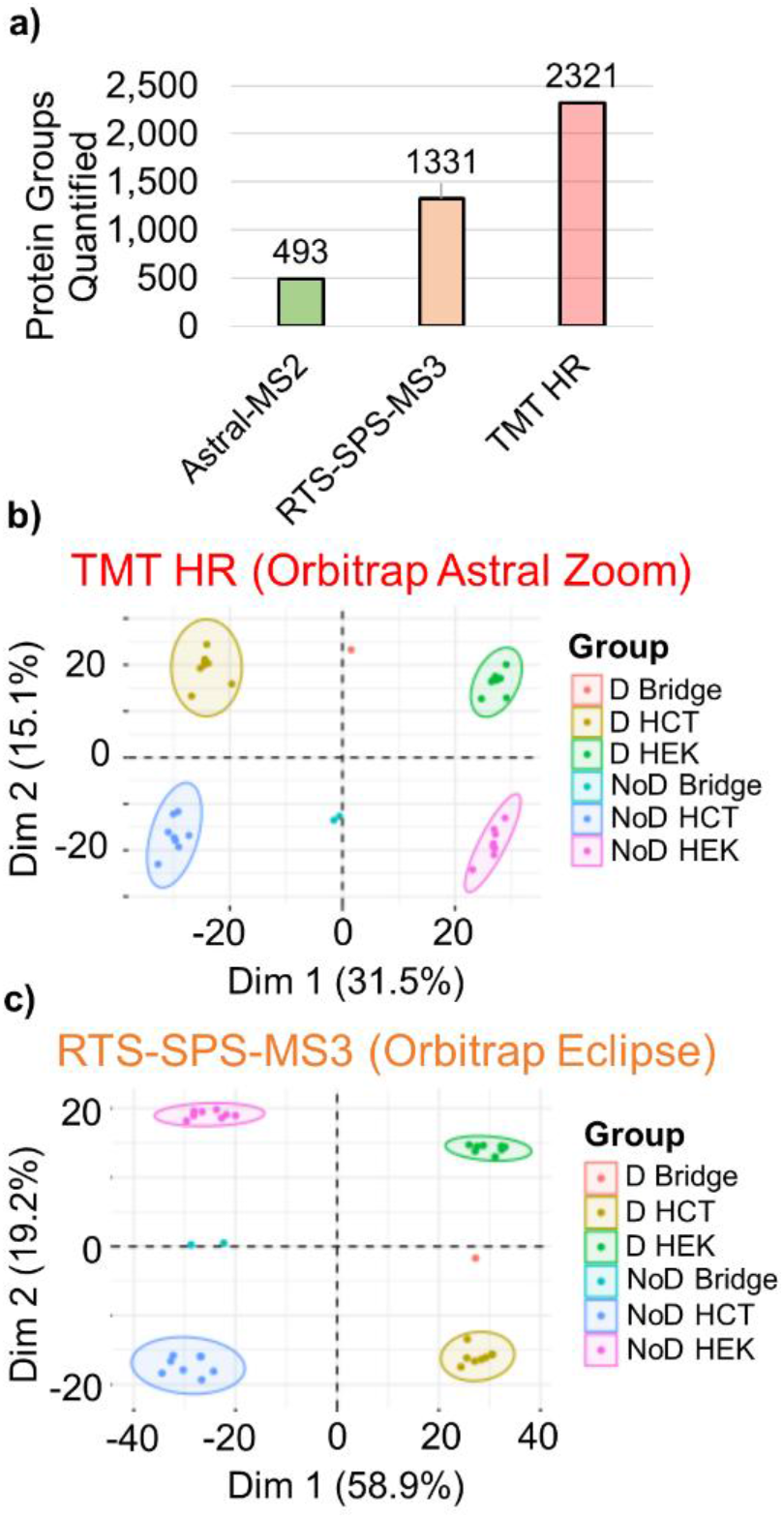
a) Comparison of proteins quantified in 90-minute analysis of 250 ng 2-cell line 35-plex sample via standard Orbitrap Astral Zoom MS2, Orbitrap Eclipse RTS-SPS-MS3 and Orbitrap Astral Zoom TMT HR mode. b) Principle component analysis of TMT HR and c) RTS-SPS-MS3 protein quantities in each channel.

Orbitrap MS3 analysis necessarily requires long acquisition cycles and ion accumulation times, both for the ion losses inherent to MS3 and the measurement transient to resolve the reporter quadruplets. Real-time search and synchronous precursor selection serve to compensate for these limitations. What the MS3 method excels at, however, is in the reduction of chemical interferences from co-isolated labeled peptides, and thus returning reliable quantities. As shown in Figure 8b–c, the tight clustering observed in principle component analysis (PCA) qualitatively suggests similar quantitative precision between the TMT HR and RTS-SPS-MS3 methods, although with TMT HR, the deuterium and cell line effects (31.5% + 15.1%) account for a smaller proportion of the variance than with RTS-SPS-MS3 (58.9% + 19.2%). Strikingly, the deuterium effect accounts for 58.9% of variance (PC1) with RTS-SPS-MS3, over 3x more than the 15.1% (PC2) with TMT HR; since the deuterium effect arises from slightly different retention times between deuterated and non-deuterated peptides, this reduction in variance may arise from the superior speed of the Orbitrap Astral Zoom mass spectrometer.

The Orbitrap Astral Zoom mass spectrometer is not capable of MS3 or synchronous precursor selection because it lacks an ion trap capable of multi-notch isolation, and its control software does not currently implement real-time search strategies to skip uninformative TMT HR quantitation scans. In this regard, a more complete comparison between Orbitrap and Astral analyzer in TMT HR mode cannot be made, though it is reasonable to assume that either or both features would aid performance.

One important application of labeled peptide analysis not addressed in this investigation is the use of multiplexing to pool low-input samples, such as single cells, thereby amplifying the number of ions available for precursor detection and peptide identification.

It is reassuring that a method such as multi-pass operation has, after considerable development, found a valuable application. Multi-pass operation is very superficially attractive to an analyzer designer for its resolving power, but in most popular applications is rendered useless by ion losses, mass range restrictions, and hampered by other difficulties with tuning and calibration. Beyond niche targeted applications where higher resolution is required, there remain other prospects for the method which are as yet undeveloped, including perhaps construction of high-resolution, high-dynamic range MS1 survey scans out of multiple measurements at differing mass ranges, and possibly the measurement of TMTc complemented ions^22^.

## CONCLUSION

The use of multi-pass operation, or TMT HR mode, to improve Astral analyzer resolving power, has been shown to sufficiently resolve 3 mDa TMTpro reporter ion multiplets and unlock the capability to quantify TMTpro 32- and 35-plex labeled samples. The TMT HR method, introduced to the Orbitrap Astral Zoom mass spectrometer, with Orbitrap survey scans, parallel Astral MS2 identification of labeled peptides, and paired multi-pass MS2 reporter ion quantification scans, is demonstrated to provide effective analysis of complex 32- and 35-plex labeled samples, without substantial sacrifice of analytical depth. It is anticipated that the near doubling of experimental throughput this method unlocks will prove valuable to proteomic studies, while low-load and single cell investigation will benefit from combining additional multiplexed samples with the highly sensitive Astral analyzer.

## ASSOCIATED CONTENT

### Supporting Information

Additional details of automated multi-pass mode calibration, multiplexed HeLa data analysis, experimental method editor and data processing inputs (PDF).

## AUTHOR INFORMATION

### Author Contributions

The manuscript was written through contributions of all authors. All authors have given approval to the final version of the manuscript.

### Notes

The authors declare the following competing financial interest(s): HS, CR, JK, MZ, TNA, BH, ED, RO, DG, JP, DN, PC, FB, BD, MW, AW, RB, DCF, ED, AM, VZ and CH are employees of, and SPG a consultant for, Thermo Fisher Scientific, the manufacturer of instrumentation used in this research.

## ACKNOWLEDGMENT

The authors would like to acknowledge the advice and support of colleagues throughout Thermo Fisher Scientific, as well as academic and industrial collaborators.

