## Supporting Information for "High Resolution Multi-Pass Astral Analyzer Quantification Enables Highly Multiplexed 35-Plex Tandem Mass Tag Proteomics"

Thermo Fisher Scientific, 11 Hannah-Kunath Str., 28199 Bremen, Germany.

[†]Department of Cell Biology, Harvard Medical School, 240 Longwood Ave., Boston, MA 02115, United States.

[ǂ]Thermo Fisher Scientific, 3747 N Meridian Rd, Rockford, IL 61101, United States.

[‡]Thermo Fisher Scientific, 355 River Oaks Pkwy, San Jose, CA 95134, United States.

### SUPPORTING INFORMATION

**Automatic tuning of the TMT HR Mode:** The relevant analyzer voltages of the TMT HR mode depend on each other and thus span a multi-dimensional parameter space. An automatic tuning routine was implemented where a genetic search algorithm is utilized to first maximize the transmission and suppress overtones, low-intensity artifacts caused by ions travelling at a different number of oscillations than the principal peak, by finding the optimal ion foil and relay prism voltages (injection and trapping mode) as depicted in Supplemental Figure 1a. In a second step, a linear scan of the focusing mirror electrode voltage optimizes the resolution over a range of ion targets (Supplemental Figure 1b). One may see that the peak broadens terribly without this focus compensation voltage, or with a positive potential, and comes into focus around -25 to -30 V.

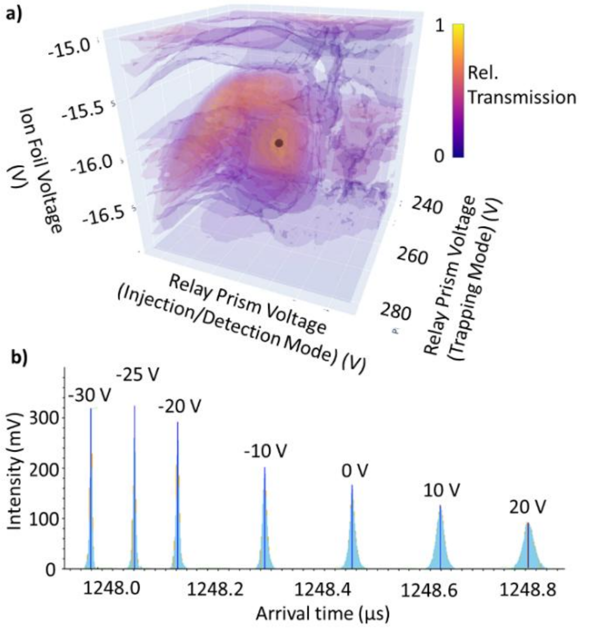


Supplemental Figure 1. a) 3-dimensional representation of the relative ion transmission. A genetic algorithm is used to find the optimal analyzer voltages for the relay prism (injection and trapping mode) and the ion foil (found optimum marked by dark dot). b) Superimposed profile spectra showing how temporal focus of the m/z 138 ion bunch is shifted to the detector plane by applying a correction voltage to the first electrode of the ion mirrors.

**Additional HeLa 32-plex analysis**: Further results of analysis of a 1:4 ratio 32-plex HeLa sample by standard Orbitrap Astral TMT quantification and via the TMT HR method are shown in Supplemental Figure 2 below. Attempting to apply the standard Astral TMT analysis method to a 32-plex sample returned 10 quantified proteins, vs 1744 via TMT-HR mode, as the inability to resolve reporter ion channels leads to an almost complete number of rejected spectra with apparently empty channels. The distribution of reporter full width half maximum resolving power could be seen to increase from a median of 70k to well over 100k with the use of TMT-HR mode, while missing channels in standard Astral MS2 TMT quantification are shown creating a spike at R=0.

Notably there remain some TMT HR mode reporter ion peaks with apparently relatively low and very high resolution. Some of the lower resolution peaks will be a real consequence of intense space charge, though perhaps much of the spread is simply due to jitter associated with measuring low-lying peaks. Single ion peaks also are very narrow, and can show apparent resolving power of 300k.

Supplemental Figure 2c plots the intensity ratio of two reporter channels, the 128CD vs 128ND, which should reflect the deliberate 4:1 concentration ratio they were made up with, against peak resolution. This fortunately is well matched, on average, with a reasonably narrow distribution that interestingly enough has little visible relation to resolution. There was little evidence of artifacts or heavy peak coalescence at higher intensity, though this is likely obscured by both the low intensity of peaks in this experiment, and the nature of the test sample where 4x intensity channels largely neighbored 1x intensity channels, limiting the impact of overlap.

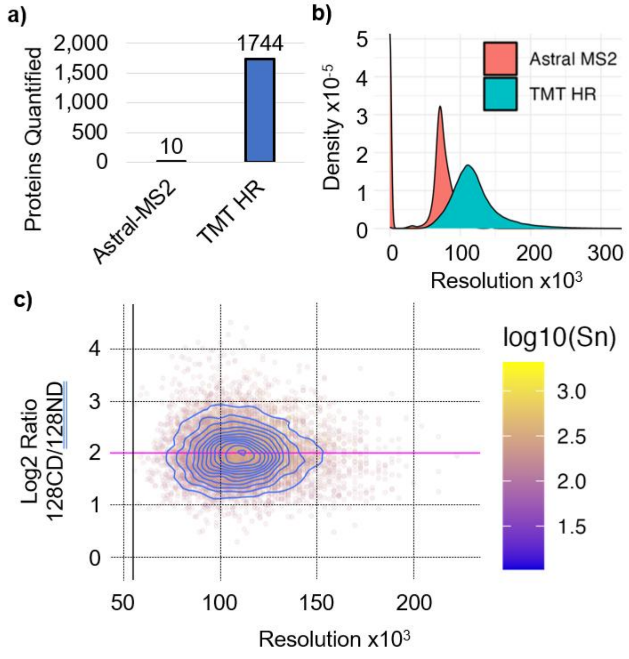


Supplemental Figure 2. a) Comparison of proteins quantified from 32-plex Hela sample in standard Orbitrap Astral TMT analysis and with TMT HR mode. b) Distribution of reporter ion peak resolution measured in standard and TMT HR modes. c) Distribution of measured ratio of 128CD to 128ND channels (theoretical 4:1 ratio) vs peak resolution.

**HeLa 35-plex analysis:** Supplemental Figure 3 shows further experiments carried out at a later date, with a 1:4 ratio 35-plex labelled sample rather than the previous 32-plex, during the validation process of the Orbitrap Astral Zoom mass spectrometer. In this case an 88-minute gradient was used and the amount of TMT labelled HeLa increased from 500 to 1000 and 1500 ng. Higher loads show a moderate improvement in number of quantified proteins and peptides.

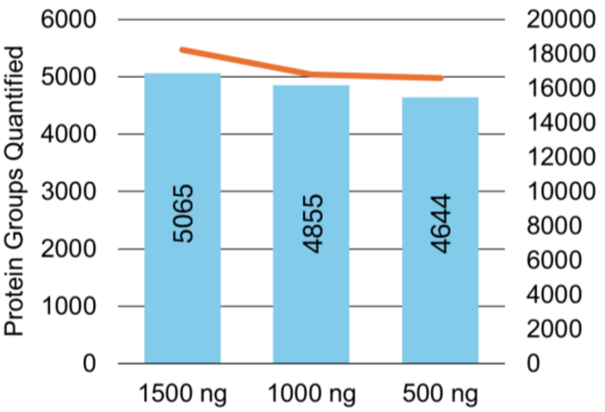


Supplemental Figure 3. Proteins and peptides quantified from 88-minute gradient analyses of TMT labelled HeLa.

**Additional method and processing details:** Supplemental Figure 4 shows method editor inputs for a typical method incorporating reporter ion quantification via TMT HR mode, as an aid for reproduction or adaptation. Similarly, Supplemental Figure 5 shows several Proteome Discoverer 3.3 workflow and filter options for data analysis. Notably there are filter options for deviant reporter ion peaks, to detect peaks likely to suffer either insufficient ion number to quantify on, or an excessive ion number that would trigger overlap and crosstalk with neighbouring channels. At present if a single reporter channel triggers the signal to noise or resolution filter (set to 0 and not used in the method of Supplemental Figure 5), the entire spectrum is dropped, which is perhaps somewhat wasteful to the other properly quantified channels, if robust.

| No | Time | Duration [min] | Flow [µl/min] | %B | Volume [µl] | No. of Column Volumes |
|---|---|---|---|---|---|---|
| 1 | 0.000 | Run | | | | |
| 2 | 0.000 | 0.000 | 0.500 | 5.0 | 0.00 | 0.00 |
| 3 | 1.000 | 1.000 | 0.500 | 5.0 | 0.50 | 0.28 |
| 4 | 1.100 | 0.100 | 0.300 | 7.0 | 0.04 | 0.02 |
| 5 | 59.100 | 58.000 | 0.300 | 36.0 | 17.40 | 9.80 |
| 6 | 79.100 | 20.000 | 0.300 | 42.0 | 6.00 | 3.38 |
| 7 | 79.100 | Column Wash | | | | |
| 8 | 80.000 | 0.900 | 0.300 | 90.0 | 0.27 | 0.15 |
| 9 | 87.000 | 7.000 | 0.300 | 90.0 | 2.10 | 1.18 |
| 10 | 87.000 | Stop Run | | | | |
| 11 | 87.000 | Column Equilibration | | | | |

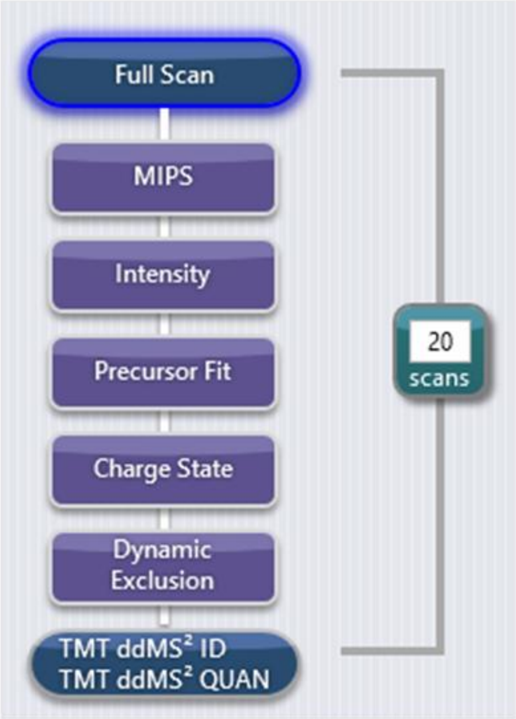


**Theoretical Precursor Envelope Fit Properties**

| | |
|---|---|
| Fit Threshold (%) | 70 |
| Fit Window (m/z) | 1.2 |

**TMT IDENTIFICATION SCAN**

| | |
|---|---|
| Multiplex Ions | ☐ |
| Isolation Window (m/z) | 1.2 |
| Collision Energy Type | Normalized |
| HCD Collision Energy (%) | 32 |
| Detector Type | Astral |
| Scan Range Mode | Define m/z Range |
| Scan Range (m/z) | 150-2000 |
| Normalized AGC Target (%) | 100 |
| Maximum Injection Time (ms) | 5 |

**TMT QUANTIFICATION SCAN**

| | |
|---|---|
| TMT HR Mode | ☑ |
| Isolation Window (m/z) | 0.5 |
| HCD Collision Energy (%) | 55 |
| AGC Target | Custom |
| Normalized AGC Target (%) | 300 |
| Maximum Injection Time (ms) | 50 |

Supplemental Figure 4. Method editor details for TMT HR MS method and LC gradient.

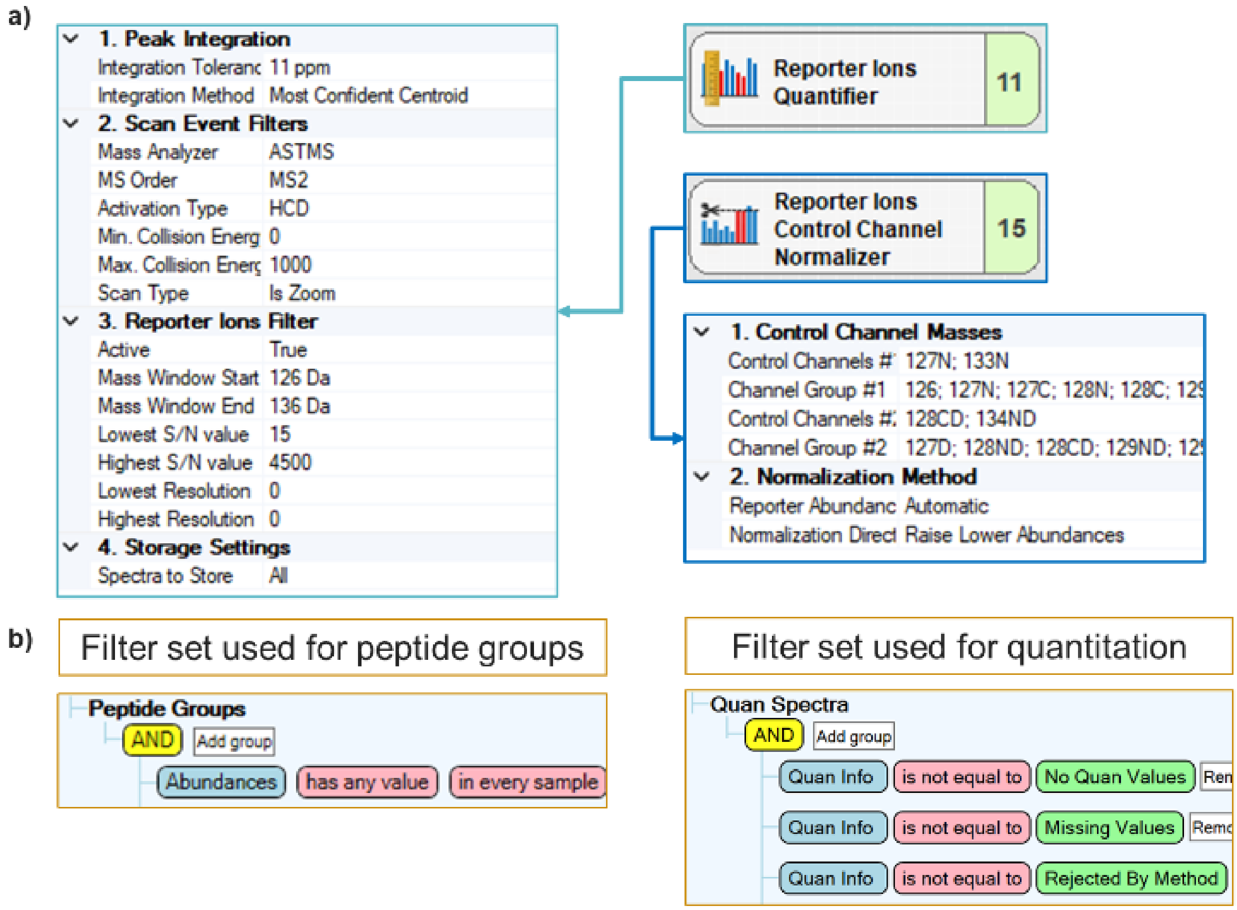


Supplemental Figure 5. a) Workflow and b) filter set in Proteome Discoverer 3.3 for analysis of TMT HR mode results.
